# Specific killing of Ewing sarcoma by TCR-T cells targeting public neogene-encoded antigens

**DOI:** 10.64898/2026.06.20.733160

**Authors:** Ana I. Lalanne, Céline Collin, Floriane Petit, Mégane Lacaud, Yago A. Arribas, Aurélie Darbois Delahousse, Amaury Leruste, Mariia K. Koshkina, Kyle A. Raymond, Paul Klein, Julien Vibert, Sakina Zaidi, Sandrine Grossetête, Jill Pilet, Karine Laud-Duval, Setareh Aflaki, Camille Jamet, Wolfgang Faigle, Jaxaira Maggi, Montserrat Carrascal, Silvia Menegatti, Jaime Fuentealba, Marion Alcantara, Joshua J. Waterfall, Olivier Lantz, Olivier Delattre

**Affiliations:** Laboratory of Clinical immunology, Department of Diagnostic and Theranostic Medecine, Institut Curie, Paris, France; Centre d’investigation Clinique Inserm 2501, Institut Curie, Paris, France; INSERM U1330, Diversity and Plasticity of Childhood Sarcoma Lab, PSL Research University, SIREDO Oncology Center, Institut Curie Research Center, Paris, France; INSERM U932, Immune responses and cancer Lab, PSL Research University, Institut Curie Research Center, Paris, France; CellAction, Center for Cancer Immunotherapy, INSERM U932, Institut Curie, Saint-Cloud, France; INSERM U1330, Integrative Functional Genomics of Cancer Lab, PSL Research University, Institut Curie Research Center, Paris, France; Department of Translational Research, PSL Research University, Institut Curie Research Center, Paris, France; Biological and Environmental Proteomics, Institut d’Investigacions Biomèdiques de Barcelona-CSIC, IDIBAPS, Roselló 161, 6a planta, 08036 Barcelona, Spain; CSIC/UAB Proteomics Laboratory, Universitat Autònoma de Barcelona, 08193 Cerdanyola del Vallès, Spain

**Author notes:** Corresponding authors, Dr Olivier Delattre, Dr Olivier Lantz, Dr Joshua Waterfall. equal contribution (Ana I. Lalanne, Céline Collin, Floriane Petit). shared senior authorship and corresponding authors (Joshua J. Waterfall, Olivier Lantz, Olivier Delattre).

## Abstract

EWSR1::FLI1, the oncogenic chimeric transcription factor driving Ewing sarcoma (EwS)induces expression of exquisitely EwS-specific neogenes (Ew_NGs) through neomorphic binding and transcription activation at GGAA microsatellites in genomic regions that are silent in normal tissues. We show that peptides encoded by Ew_NGs are presented on HLA-I complexes on EwS cells. The cytokine secretion of CD8+ T cells specific for Ew_NG-encoded HLA-I-bound peptides is activated by all HLA-I-matched EwS cells but not by non-EwS cells. These T cells kill EwS cells in an HLA-I restricted manner. This cytotoxicity is dependent on the expression of EWSR1::FLI1 and of the corresponding Ew_NG. It can be reproduced by transduction of the TCR into donor T cells (TCR-T) which kill EwS cells in vivo. Moreover, we show that neither off target nor allogeneic activation are observed with TCR-T thus paving the way for cell therapy in relapsed/resistant EwS patients for which therapeutic options are very limited.

**Statement of significance:** The chimeric transcription factor EWSR1::FLI1 generates tumor-specific neogenes encoding neoantigen presented by the HLA-I molecules of Ewing cells. Neoantigen-specific CD8^+^ T-cell clones and engineered TCR-T cells can selectively recognize and kill EwS tumor cells *in vitro* and *in vivo*.

## Introduction

Ewing sarcoma (EwS) is the second most frequent bone and soft tissue cancer of childhood and adolescence. EwS is a highly aggressive cancer with a 5-year overall survival of only 30% for patients with metastases. Metastatic disease is often resistant to intensive therapy and associated with acute and chronic adverse effects. EwS possesses one of the lowest mutation rates among cancer, does not typically exhibit high T cell infiltration, and does not usually respond to monotherapy immune checkpoint blockade (1–5). However, preliminary results from the Ewing cell vaccination Vigil study suggest that Ewing cells can generate an immune response to yet uncharacterized tumour antigens (6).

EwS is characterized by specific gene fusions between members of FET family of RNA-binding proteins and the ETS (E-twenty-six) family of transcription factors, the most frequent fusion being between *EWSR1* and *FLI1* in 85% of cases (7,8). EWSR1::FLI1 acts as an aberrant tumour-specific transcription factor reprogramming the genome through binding DNA at two types of DNA motifs, i.e. bona fide ETS family sites centred on a single GGAA/T motif and GGAA microsatellite repeats. As compared to wild-type ETS transcription factors, EWSR1::FLI1 has the neomorphic activity to generate neo-enhancers upon binding GGAA microsatellite sequences (8).

We recently showed that EWSR1::FLI1 can drive transcription and processing of a specific set of novel spliced and polyadenylated transcripts within otherwise transcriptionally silent regions of the genome upon binding GGAA microsatellite sequences (9). These novel transcribed genomic regions, that are not expressed in any normal tissues, were termed Ewing-specific neogenes (*Ew_NG*). We further used both ribo-seq and whole cell proteomics to demonstrate that some of these neogenes are translated (9). The exquisite tumour specificity of these neogenes and their highly reproducible expression across EwS patients make them attractive as a source of neo-epitopes.

We investigate here the possibility that open reading frames (ORFs) derived from these neogenes represent a public source of immunogenic tumor-specific neoantigens. To explore this hypothesis, we performed ribosome profiling and HLA-I immunopeptidomics on EwS cell lines and patients-derived xenografts (PDXs) models. A set of neopeptides derived from *Ew_NGs* was specifically identified in EwS samples. We found that tumour-specific peptides are presented to T cells by Ewing cell HLA-I molecules and functionally activate T cells through HLA-restricted recognition. These T-cells, as well as engineered TCR-T, kill Ewing cells *in vitro* and *in vivo*. Our findings thus pave the way for antigen specific immunotherapies in Ewing sarcoma.

## Results

### *Ew_NGs*-encoded peptides contribute to the EwS-specific HLA-I immunopeptidome

To investigate whether peptides encoded by these *EWSR1::FLI1*-specific neogenes (*Ew_NGs*) can be processed and presented by HLA-I molecules on the surface of Ewing cells, we performed immunoprecipitation of HLA-I molecules in five Ewing cell lines and five PDX followed by analysis of the eluted peptides using LC-MS/MS. We thus generated a database of EwS immunopeptidomics MS spectra. Two search tools, Proteome Discoverer and MSFragger, were used to identify matches with the human reference proteome and with a database of *Ew_NGs* (9). Proteome Discoverer identified 25 *Ew_NGs*-derived peptides, 17 of which being compliant with the HLA-I alleles of the corresponding sample. MSFragger identified 22 *Ew_NG*-derived peptides, with 17 being HLA-I compliant. Sixteen peptides were common to both analyses (Fig. 1A and B, Table 1, Supplementary Table S1).

**Fig. 1:**
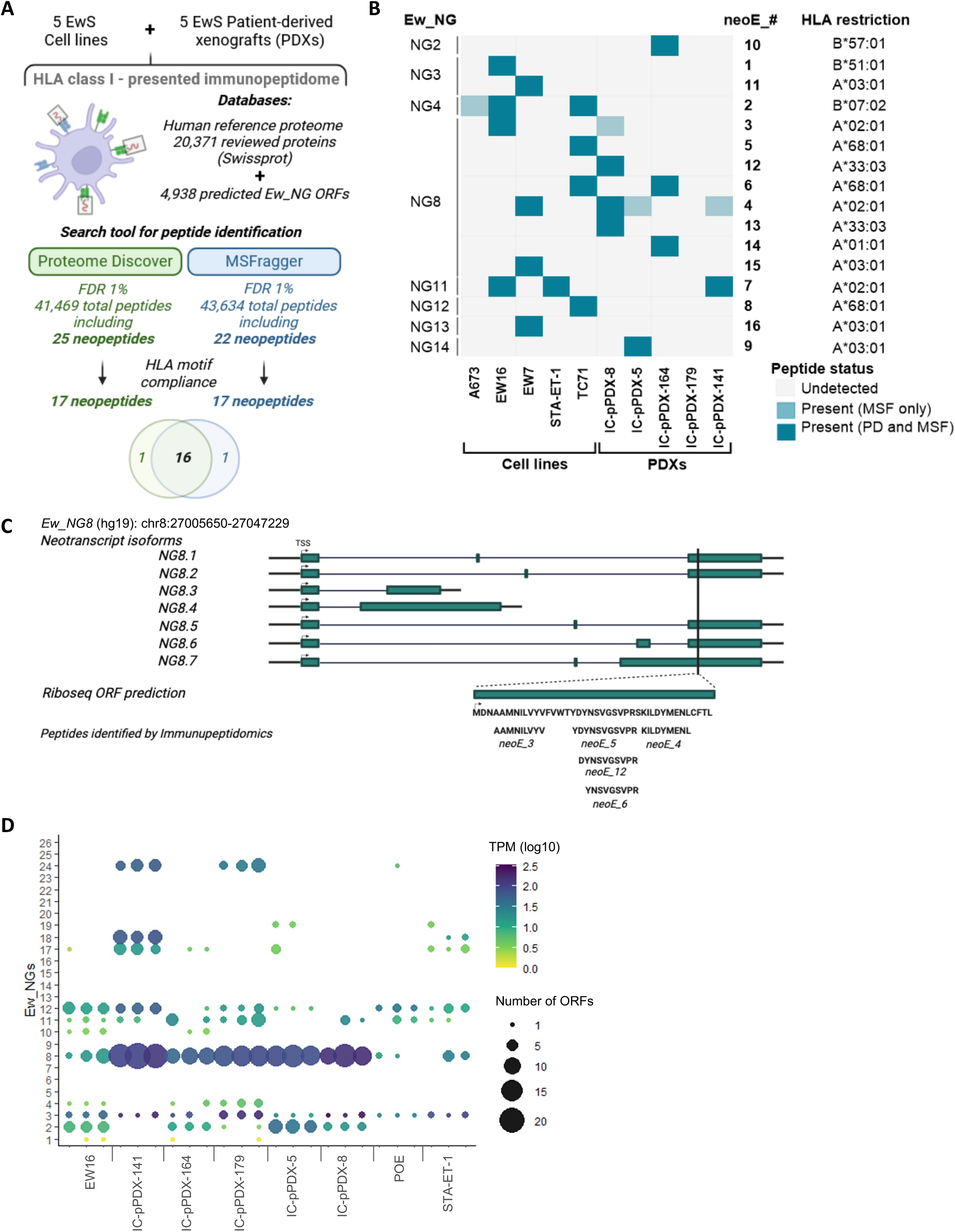
Identification of MHC-I bound peptides encoded by the EwS neogenes. (A) Workflow for the identification of EwS neogene-derived peptides by MS-based immunopeptidomics. (B) Summary of EwS neogene-derived peptides found by immunopeptidomics in each EwS sample analyzed. (C) Genomic location of ribosome protected fragments detected by Ribo-seq and peptides detected by immunopeptidomics for *Ew_NG8*. (D) Expression of *Ew-NGs*-derived RPFs. Bubble color represents mean expression level of *Ew_NG*-specific RPFs in all EwS cell lines and PDXs. Dots size shows number of detected ORFs for each *Ew_NGs* in all tested samples. Cartoons created using Biorender.

**Table 1.**
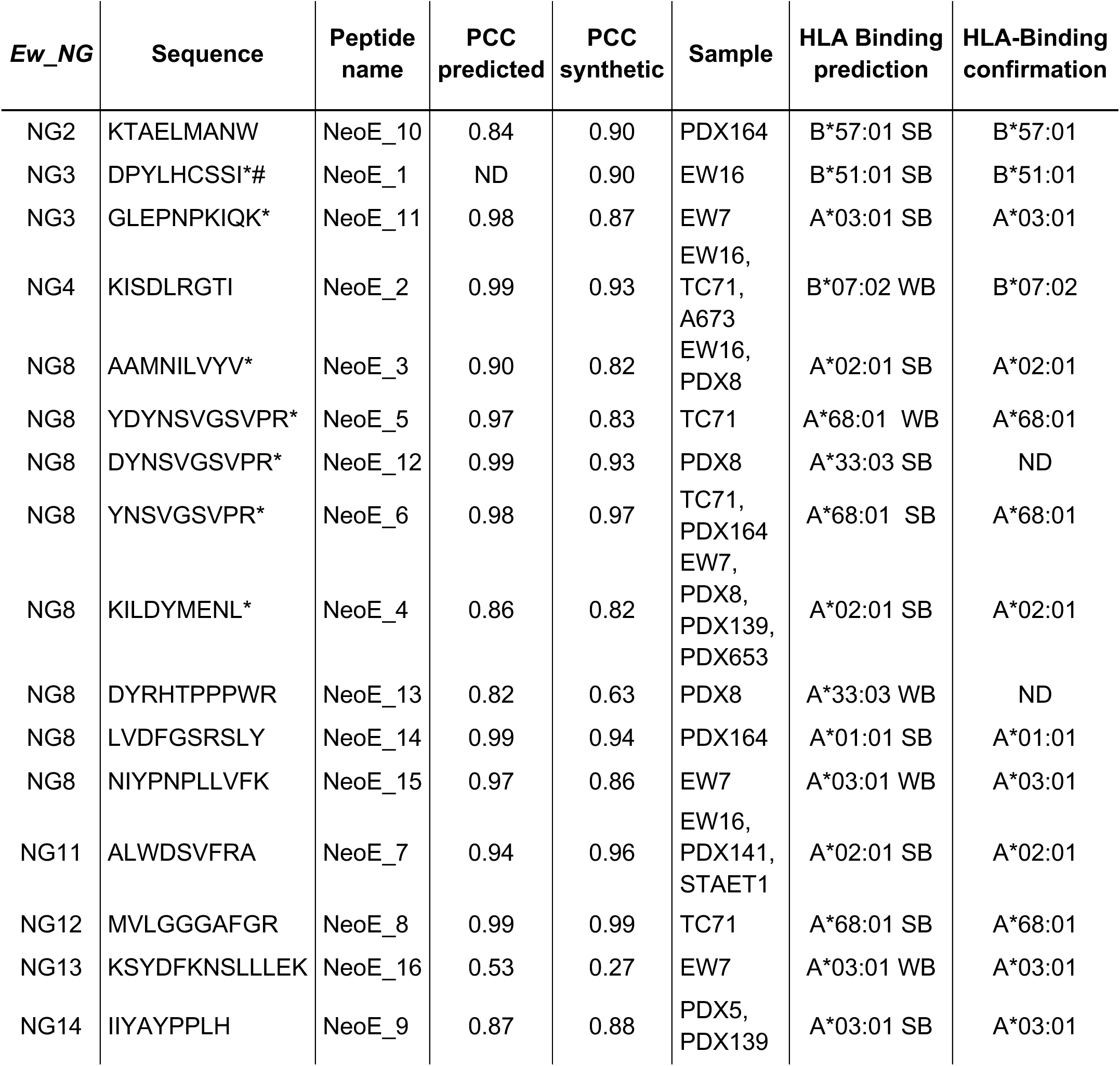
*Ew_NG*-encoded peptides identified by immunopeptidomics. *, Part of an ORF predicted by Ribosome profiling; #, this gene corresponds to an expressed sequenced tag (EST) identified in Ewing cells (hCG1821234); ND, Not Done; PCC, Pearson Coefficient Correlation; NetMHCPan 4.0 predictions: Strong Binder (SB) Rank lower than 0.05; Weak Binder (WB) between 0.05 and 5.

The *Ew_NG*-encoded peptides length distribution peaked at 9 amino acids consistent with the expected length for HLA-I presented peptides (Table 1) and similar to that of peptides from annotated genes (Supplementary Fig. S1A). Additionally, for both *Ew_NG*-derived peptides and annotated peptides the ratio of calculated hydrophobicity to measured HPLC retention time was similar (Supplementary Fig. S1B). The robustness of *Ew_NG*-derived peptide detection was further corroborated by similar MS/MS spectrum scores with annotated peptides (Supplementary Fig. S1C).

To further validate EwS peptide detection, we compared immunopeptidomics MS/MS spectra to corresponding predicted and synthetic peptide spectra (Supplementary Fig. S1D and E, Table 1). Several peptides were found in multiple samples with identical HLA-I restriction (Fig. 1B, Supplementary Table S2). The sixteen peptides were encoded by eight distinct *Ew_NGs* (Fig. 1B, Supplementary Table S3). *Ew_NG8* was the most represented, with a total of eight peptides encoded by its various isoforms. Five of these isoforms include an ORF encoding five HLA-I presented peptides (Fig. 1C). Further reinforcing the Ewing specificity of these peptides, none of them were detected across 407 immunopeptidomic samples from normal tissues available in public databases (10).

We compared immunopeptidomic results with Ribosome profiling from the same samples. In addition to the two previously analyzed cell lines (9), we also investigated the three additional EwS cell lines and the five EwS PDX models analyzed by immunopeptidomics. Following removal of rRNA and tRNA, ribosome-protected fragments (RPFs) were aligned with a database comprising the Gencode v19 transcriptome and *Ew_NGs*, enabling the assignment of RPFs to the EwS transcriptome (Overview in Supplementary Fig. S2A). As expected, the mean length of *Ew_NG* ORFs with RPFs was shorter than that of annotated genes (Supplementary Fig. S2B). The quality of the RiboSeq data was validated by observing a trinucleotide periodicity consistent with the known reading frames of coding genes (Supplementary Fig. S2C). Only ORFs detected in all three replicates of a given sample were retained. The detected ORFs were classified into six categories: canonical (coding sequences), extension, readthrough, upstream (uORF), downstream (dORF) or *Ew_NG* (Supplementary Fig. S2D).

Further processing of the 663 *Ew_NG* ORFs detected by ribosome profiling, by grouping overlapping ORFs and filtering to retain the most 5’ ones, yielded 91 unique *Ew_NG* ORFs associated with RPFs corresponding to 12 *Ew_NGs* (Figure 1D, Supplementary Fig. S2E). The ORFs detected by RiboSeq on *Ew_NG8* included all five peptides found by immunopeptidomics MS/MS. They had multiple potential start codons including methionines (Fig. 1C). The ribosome protection of this ORF was observed in most EwS samples with some heterogeneity in abundance (Fig. 1D). Similarly, the number of ORFs and their TPM values for other *Ew_NGs* varied across samples (Fig. 1D; Supplementary Fig. S2E). As indicated above, the sixteen peptides found by immunopeptidomics were encoded by 11 distinct *Ew_NGs* ORFs from 8 *Ew_NGs*. Riboseq data indicates that two of these (*Ew_NG 3* and *8)* were unambiguously associated with RPFs detected by Riboseq (Supplementary Fig. S2F). The consistency of the findings between two orthogonal approaches, Riboseq and immunopeptidomics, and the number of detected peptides highlights *Ew_NG8* as the strongest candidate for subsequent experiments. Sensitivity issues of both approaches may account for the only partial overlap between Riboseq and immunopeptidomics data.

Altogether, this data indicates that the chimeric transcription factor EWSR1::FLI1 induces the expression of *EwS_NGs* that are further translated into *Ew_NG*-derived neoproteins which are subsequently processed into neopeptides and presented by HLA-I molecules.

### Identification of CD8^+^ T cells specific for *Ew_NG*-encoded peptides

Next, we addressed whether the *Ew_NG*-encoded peptides presented by HLA-I molecules can be detected by CD8^+^ T cells and generate effective anti-Ewing cell responses. First, we confirmed peptide binding predictions to HLA-I molecules using an HLA monomer refolding assay. The HLA-I complex stabilization by the candidate peptides was compared to control peptides. All fourteen tested EwS peptides stabilized (more than 50% of control) at least one HLA-I allele of the source EwS samples (Fig. 2A).

**Fig. 2:**
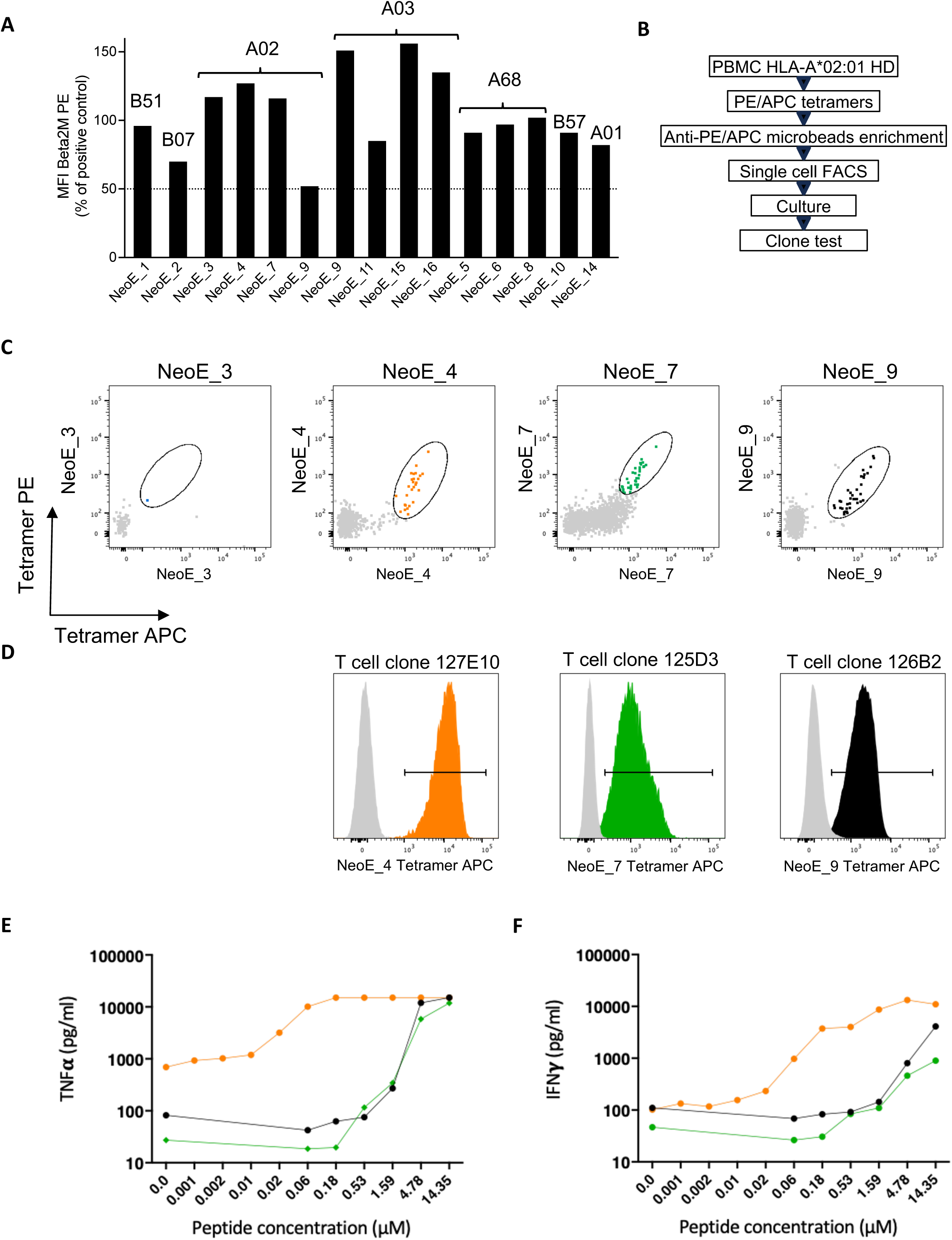
Identification of T cells specific for Ew_NG-derived peptides. (A) HLA-I in vitro monomer binding assay of Ew_NG-derived peptides. Relative binding compared to positive peptide controls is displayed for each HLA. Above 50% of binding relative to positive control is considered good binding. (B) Workflow for the identification of T cell clones against Ew_NG-derived peptides. (C) Tetramer-positive cells among CD3/CD8 enriched T cells for Ew_NG-derived peptides. (D) Clonal expansion and purity analysis using tetramers loaded with peptide NeoE_4 (orange), NeoE_7 (green) and NeoE_9 (black). Grey histograms correspond to an unrelated CD8+ T cell clone stained with the same tetramer. (E, F) (E) TNFα and (F) IFNγ cytokine secretion after stimulation of T cell clones with increasing concentration of the peptide NeoE_4 (orange), NeoE_7 (green) and NeoE_9 (black) loaded on K562-HLA-A*02:01 cells.

We focused on the four peptides able to strongly bind HLA-A*02:01, the most frequent allele in populations of European descent. We generated HLA-A*02:01 tetramers by associating peptides with biotinylated HLA and fluorescent streptavidin. These tetramers were used as tools to detect peptide-specific T cells from PBMCs of HLA-A*02:01+ healthy donors, using leukapheresis products to anticipate the low frequency of specific T cells in the naive pool. (11).

After staining with tetramers combined with PE and APC, CD8^+^ T cells recognizing the peptides were enriched with anti-PE and APC microbeads and magnetic columns (Fig. 2B). Tetramer-positive CD8^+^ T cells were isolated by fluorescence-activated cell sorting (FACS) as single cells on feeder cells with anti-CD3 stimulation and IL-2 and expanded to obtain CD8^+^ T cell clones (Fig. 2C).

We successfully cloned CD8^+^ T cells specific for NeoE_4 (clone 127E10), NeoE_7 (clone 125D3) and NeoE_9 (clone 126B2). These clones were specifically stained with the corresponding tetramers (Fig. 2D) and were activated by the cognate peptides loaded on the K562-HLA-A*02:01 antigen presenting cell line, resulting in the secretion of TNF-α and IFN-γ (Figure 2E-F). When comparing the functional avidity of the three clones, clone 127E10 demonstrated the highest sensitivity, detecting the lowest amount of peptide (around 20 nM). This clone was thus selected for further analysis. These data show that a naïve immune system can recognize and be activated by peptides encoded by *Ew_NGs*.

### T cell clones are activated by and kill Ewing sarcoma cells

To determine whether endogenously processed and presented NeoE_4 peptide is recognized by the 127E10 CD8^+^ T cell clone, we utilized a panel of HLA-A*02:01 positive and negative EwS and non EwS cell lines (Supplementary Fig. S3A). Each cell line was co-cultured with clone 127E10 T cells and tumor cell killing was monitored with an Annexin V assay using the Incucyte system. Very strikingly, 127E10 CD8^+^ T cells induced very rapide killing of EwS cells. Presentation of the peptide NeoE_4 by HLA-A*02:01 molecules was required, as clone 127E10 efficiently killed all five EwS cell lines expressing HLA-A*02:01 but none of the three EwS cell lines lacking this allele (Fig. 3A). Moreover, the absence of killing of non-EwS cell lines regardless of HLA-A*02:01 status (Fig. 3B) was consistent with the exclusive expression of *Ew_NG8* in EwS cells (9). Finally, blocking HLA-I presentation with a specific antibody completely abrogated EwS cell killing (Fig. 3C). Notably, none of the EwS or non EwS cell lines was killed by an anti-pp65 CMV control T cell clone generated in the same conditions (Fig. 3D). Finally, the specificity of both 127E10 and the control T cells clones was confirmed by the induction of HLA-A*02:01-positive MP41 (non-EwS) cell killing when neoE_4 or pp65 peptides were added to the coculture, respectively (Fig. 3E and F). IFN-γ secretion in the supernatant of the cell cultures was fully consistent with the killing assay (Fig. 3G and H).

**Fig. 3:**
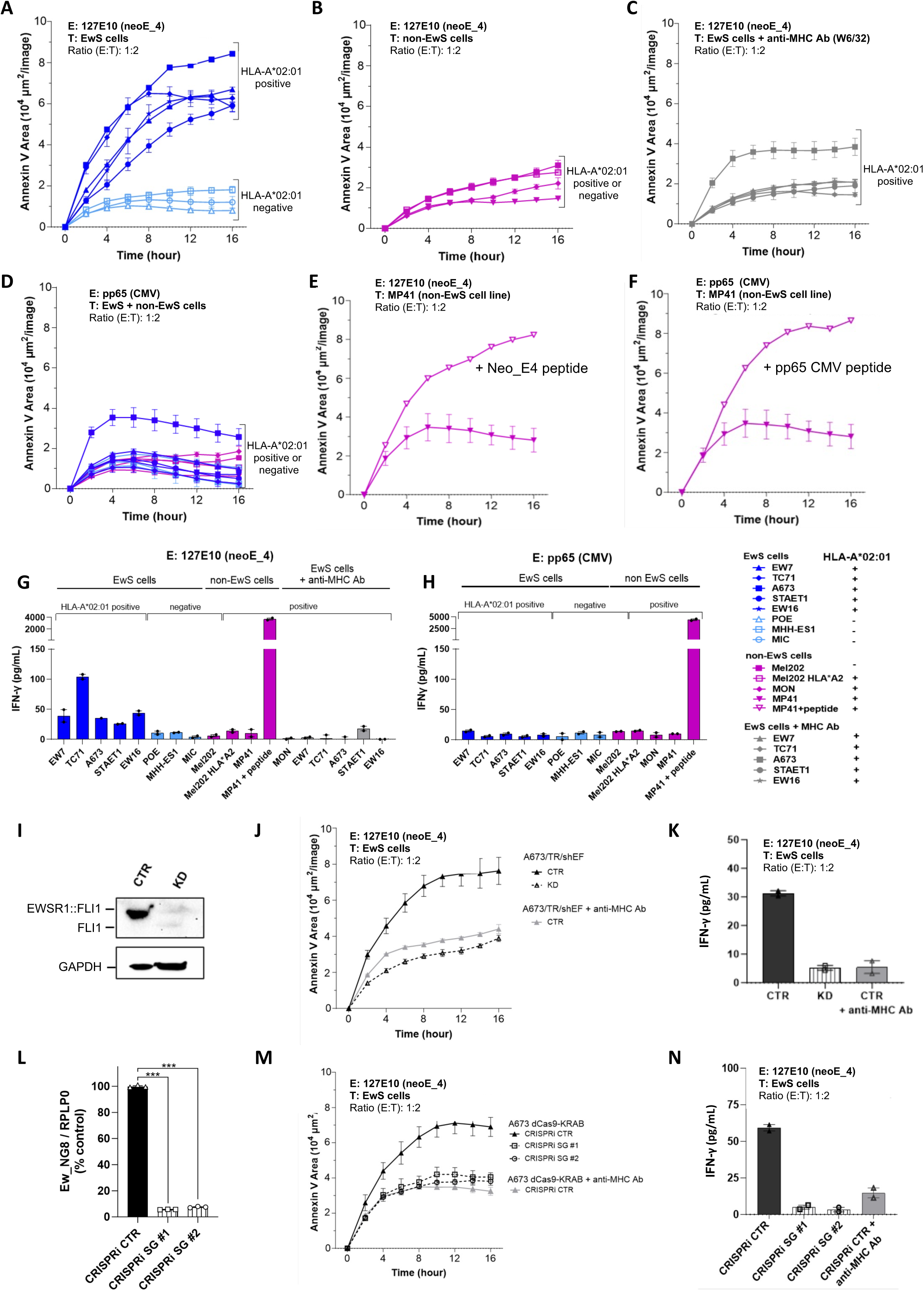
Recognition and killing of EwS cells by *Ew-NG*-derived peptides specific T Cells. (A-C) Killing assays by NeoE_4-specific 127E10 CD8+ T cells of (A) HLA-A*02:01-positive (dark blue) or HLA-A*02:01-negative (light blue) EwS cells, (B) non-EwS target cells (purple), (C) HLA-A*02:01-positive EwS cells preincubated with anti-MHC-I blocking antibody (gray). Results are represented as mean ± SEM, *n=2*. (D) Killing assay by CMV-specific CD8+ T cells of EwS cells (blue) or non-EwS cells (purple). (E,F) Killing assays by neoE_4-specific 127E10 (E) or CMV-specific pp65 T cell clones (F) of HLA-A*02:01-positive MP41 uveal melanoma cells, preincubated with ot without cognate peptides. Results are represented as mean ± SEM, *n=2*. (G, H) IFN-γ secretion of NeoE_4-specific 127E10 T cells (G) or CMV-specific pp65 T cells (H) (2.5 x 104) co-cultured with target cells (5 x 104) corresponding to an E:T ratio of 1:2. EwS cells are in blue and non-EwS cells in purple. Results are represented as mean ± SEM, *n=2*. HLA-A*02:01 restriction is indicated in the table on the right. (I) Western-Blot analysis of EWSR1-FLI1 expression in A673/TR/shEF EwS cells treated with doxycycline for 8 days to invalidate *EWSR1::FLI1* expression (KD) or untreated (CTR). (J) Killing assay and (K) cytokine secretion assay after incubation of 127E10 CD8+ T cells with the DOX-inducible A673/TR/shEF EwS cells (CTR, cells without DOX expressing *EWSR1::FLI1;* KD, after DOX induction inducing *EWSR1::FLI1*-knock down). (L) RT-qPCR of *Ew_NG8* expression in the A673 EwS cells after CRISPR interference with two different *Ew_NG8*-specific single guides (SG #1 and #2) as compared to a control sg (CTR). Results are represented as mean ± SEM, *n=3*. *** *p* value < 0.001. (M) Killing assay and (N) cytokine secretion after incubation of 127E10 CD8 T cells with A673 EwS cells upon CRISPR interference with control (CTR) or with two different *Ew_NG8*-specific single guides (sg #1 and #2). Results are represented as mean ± SEM, *n=2*. An anti-HLA-I blocking antibody is included as a control in panels J to M. E, effector cells; T, target cells.

In EwS cells, the expression of *Ew_NGs* is strictly dependent on *EWSR1::FLI1* (9). To determine whether activation of clone 127E10 was abrogated by *EWSR1::FLI1* knockdown, we used the A673/TR/shEF EwS cell line (12), in which doxycycline (DOX) induces the expression of a *EWSR1::FLI1*-specific shRNA regulating *EWSR1::FLI1* expression. Consequently, DOX addition also silences *Ew_NG8* expression which is regulated by EWSR1::FLI1 binding at a GGAA microsatellite within its promoter region (Fig. 3I and Supplementary Fig. S3B and C). Killing of A673/TR/shEF EwS cells was dramatically reduced by DOX treatment (Fig. 3J), as was IFN- γ secretion (Fig. 3K), equivalent to blocking with anti-MHC-I antibody. We previously showed that CRISPR interference targeting sequences in the vicinity of this microsatellite was sufficient to abolish expression of this neogene and did not detectably affect cell phenotypes such as proliferation or apoptosis (9). We therefore used the same two single guides (sgRNA) to silence *Ew_NG8* in the A673 cell line stably expressing a dCas9-KRAB (Fig. 3L). These guides did not influence *EWSR1::FLI1* expression (Supplementary Fig. 3D). Again, the killing was abolished by the two *Ew_NG8* sgRNAs to a level similar to preincubation of this cell line with an anti-MHC-I antibody (Fig. 3M) as was IFN-γ secretion (Fig. 3N). In contrast, a control sgRNA did not influence cell killing (Fig. 3L-N). Altogether, these data show that the killing of EwS cell lines by 127E10 T cells is strictly dependent upon the HLA-A*02:01 restriction and upon the expression of *EWSR1::FLI1* as well as of its target *Ew_NG8* which encodes the NeoE_4 peptide.

### Engineered TCR-T cells kill Ewing cells *in vitro* and *in vivo*

The next objective was to confirm that the detection of NeoE_4 on the surface of EwS cells was mediated by the T cell receptor (TCR) of clone 127E10 (TCR^127E10^). To achieve this, we sequenced the TCRs of the 127E10 clone and of the control pp65 CMV specific T cell clone. We then cloned the corresponding TCRs associated with murinized constant chains to avoid mispairing with endogenous TCRs (13) in a lentiviral vector and transduced CD8^+^ T cells from an healthy donor to obtain TCR-T cells (Fig. 4A). The TCRs indeed conferred the expected specificity to the cognate peptides, as the TCR-T cells were only stained by tetramers and activated by HLA-A*02:01 cells loaded with the specific peptides (Fig. 4B and C).

**Fig. 4:**
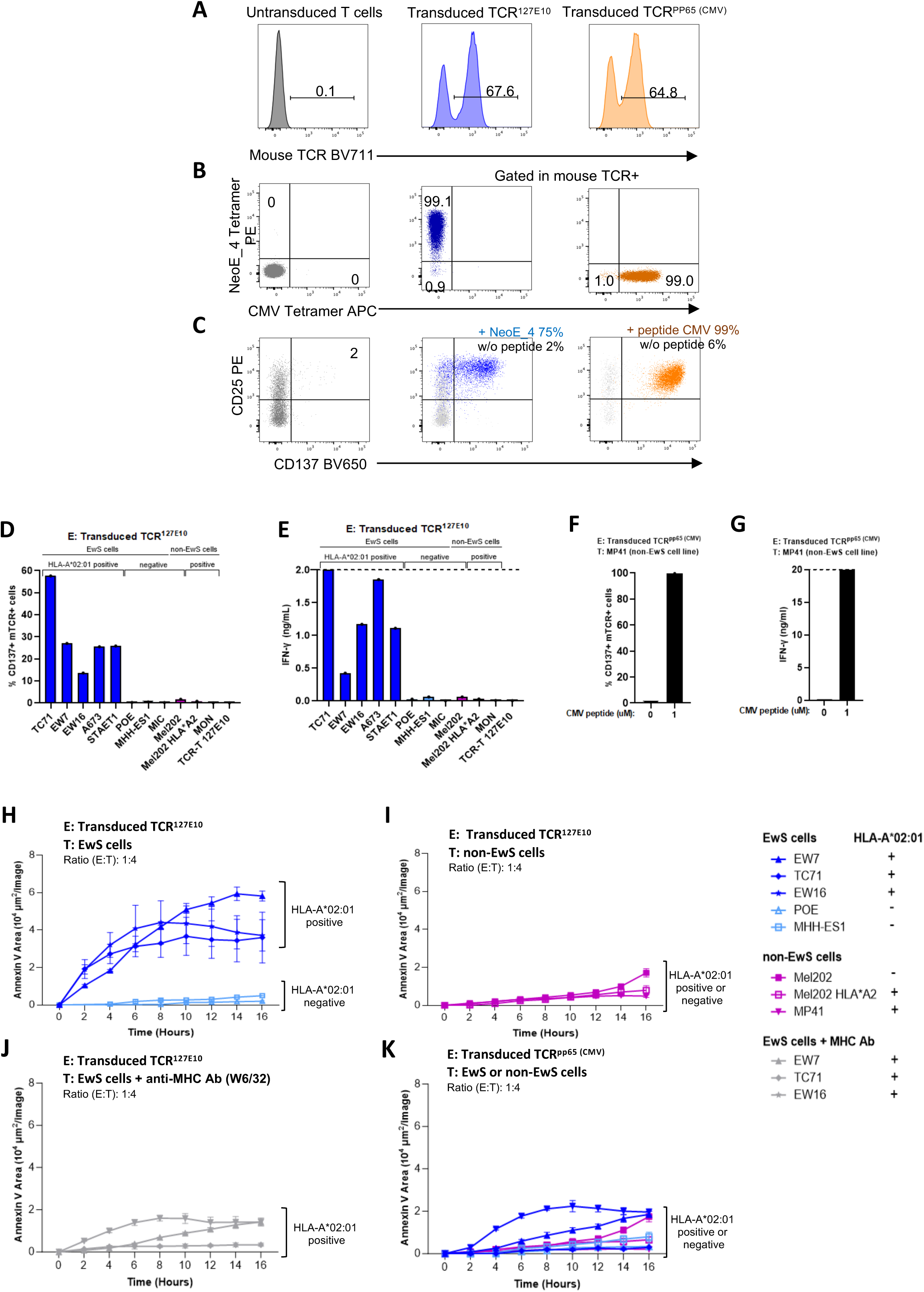
Activation and cytotoxicity of TCR127E10 transduced T cells on Ewing cells. (A) Transduction efficiency of TCR-Engineered T cells (TCR127E10, TCRpp65 or untransduced cells) represented by the frequency of mTCR-positive cells among CD8+CD3+ T cells from healthy donor (after 6 days of expansion). (B) Antigen-specific TCR-T cells were assessed by FACS analysis after tetramer staining. FACS analysis representing the frequency of expanded tetramer-positive TCR-T cells for NeoE_4 or CMV peptides. (C) Frequency of CD25+CD137+ TCR-T cells after activation with cognate peptides (blue or orange) or without peptide (gray). (D) Percentage of CD137+ TCR127E10 T cells and (E), cytokine secretion, after activation with EwS (blue) or non-EwS (purple) cells. (F) Percentage of CD137+ TCRpp65 T cells and (G), cytokine secretion after activation with MP41cells, loaded or not with the pp65 CMV peptide. Results are represented as mean ± SEM, *n=1*. (H-I) Killing assay by TCR127E10-T cells with (H), HLA-A*02:01-positive (dark blue) or HLA-A*02:01-negative (light blue) EwS cells, (I), non-EwS target cells whatever the HLA-I type (purple). (J) Similar experiments after preincubation of HLA-A*02:01 positive EwS cells with HLA-I blocking antibody (gray). (K) Killing assay by TCRpp65-T cells specific for a CMV peptide with all cell lines (EwS or non-EwS). HLA-A*02:01 restriction is indicated in the table on the right. (H-K) Results are represented as mean ± SEM, *n=2.* E, effector cells; T, target cells. All transduced TCR-T cells (12.5 x 103) were co-cultured with target cells (5 x 104) corresponding to an E:T ratio of 1:4.

We then reproduced the functional assays previously done with the T cell clones by performing co-cultures of EwS cell lines with the TCR-T cells. The capacity of tumor cell lines to activate TCR^127E10^-T cells was assessed by the expression of CD137 and by IFN-γ production (Fig. 4D-G). TCR^127E10^-T cells indeed were activated and produced IFN-γ cytokine when co-cultured with HLA-A*02:01-positive EwS cell lines but neither with the HLA-A*02:01- negative EwS cell lines nor with non-EwS cell lines (Fig. 4D and E). Neither the CMV specific TCR^pp65^-T cells nor the untransduced T cells were activated or produced IFN-γ secretion after stimulation by any cell lines (Supplementary Fig. S4B-D) unless the CMV specific peptide was added to the medium in TCR^pp65^-T cells experiments (Fig. 4F and G). Finally, and most importantly, TCR^127E10^-T cells specifically killed HLA-A*02:01^+^ EwS cell lines (Fig. 4H and I), an effect fully inhibited by an anti-HLA-I antibody (Fig. 4J). In contrast TCR^pp65^-T cells did not kill any cells in our panel (Fig. 4K). These data indicate that TCR^127E10^ is sufficient to promote *in vitro* specific killing of HLA-A*02:01-positive EwS cell lines.

To assess the potential of TCR^127E10^-T cells to eliminate Ewing sarcoma tumors *in vivo*, we generated EW7-Luc, a luciferase expressing HLA-A*02:01-positive EwS cell line. For the preparation of CD8-TCR^127E10^-T cells, CD8^+^ T cells were isolated from a healthy donor and transduced with TCR^127E10^ (Supplementary Fig. S4A). Resulting TCR^127E10^-T cells were tested *in vitro* against EW7-Luc cells using real time XCELLigence quantitative analysis and cytotoxic assay at different TCR-T/ tumor cell ratios (Supplementary Fig. S5B and C). EW7-Luc cells were injected intraperitoneally in NSG mice. Seven days later, a set of mice (n=3) was left untreated while two other sets were treated intraperitoneally with either untransduced T cells (n=6) or CD8-TCR^127E10^-T cells (n=8) (Fig. 5A). Untreated mice exhibited rapid growth of the tumor cells while untransduced T cell injection led to a small delay in tumor growth then progression in all mice but one which showed progression, probably due to some allo-reactivity (Fig. 5B). Strikingly, 7 out of the 8 mice injected with TCR^127E10^-T cells demonstrated a

**Fig. 5:**
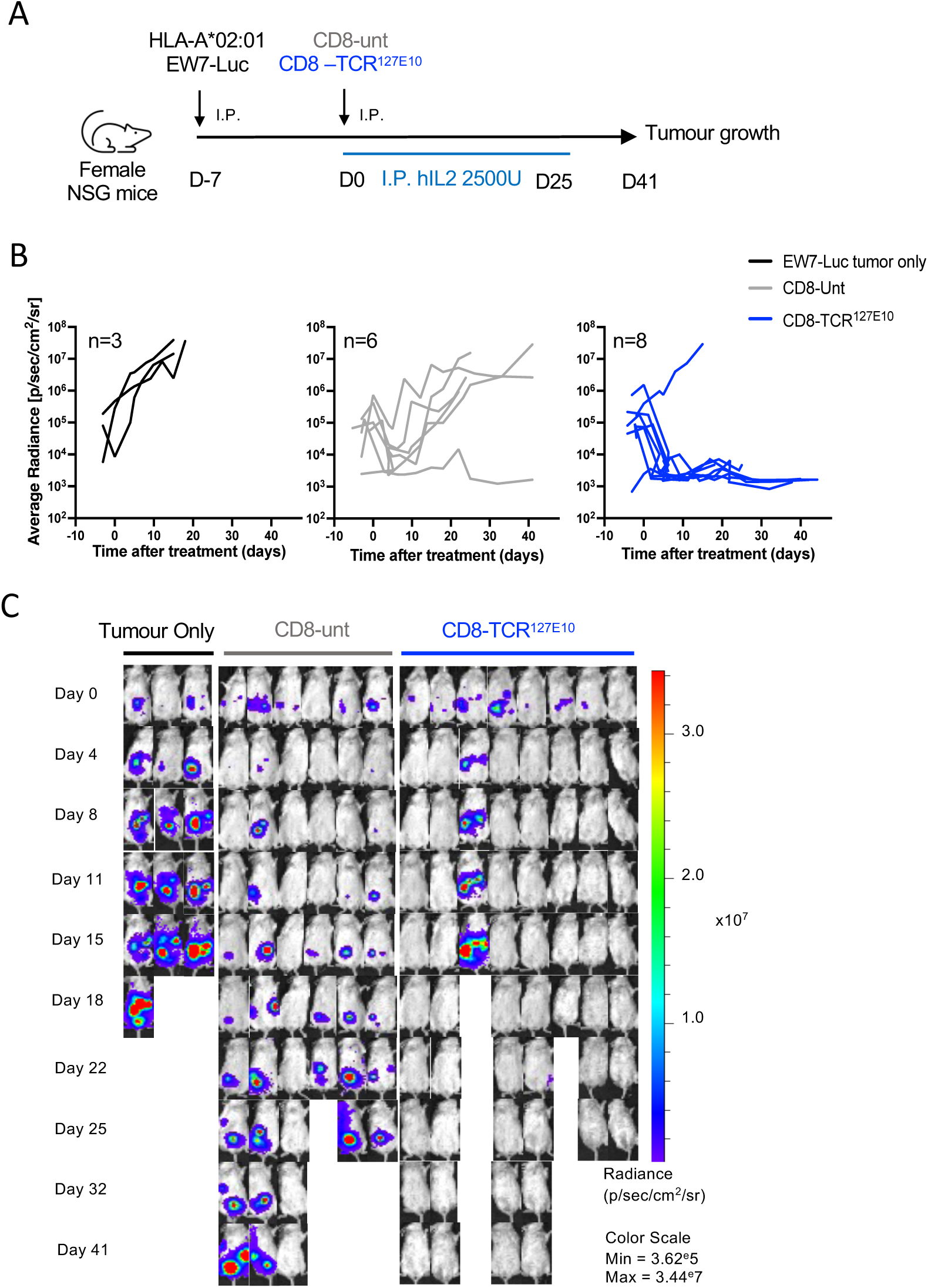
TCR^127E10^ T cells efficacy on Ewing sarcoma tumors *in vivo*. (A) Schematic overview of the protocol to assess the efficacy of the CD8^+^ TCR^127E10^ T cells *in vivo*. B) Tumour burden (average radiance, photons/sec/cm2/sr) of mice bearing HLA-A*02:01 EW7-Luc cells treated with either untransduced CD8 T cells or CD8^+^ TCR^127E10^ T cells. Radiance was measured twice per week until day 41 after T cell injection. (C) Representative bioluminescence images of each treatment group. complete regression of the tumor (Fig 5B and C). These results demonstrates that TCR^127E10^-T cells can recognize and kill Ewing cells *in vivo*.

### TCR^127E10^ does not show cross-reactivity or alloreactivity to normal peptidome

To consider TCR^127E10^-T cells for clinical use in Ewing patients, the absence of cross reactivity towards endogenous normal peptides and the absence of allogeneic activation need to be assessed. For the Ewing specificity, we applied a highly stringent method of peptide fingerprinting (14) aiming at identifying all possible off-target events. A positional scanning library of 171 peptides where each of the 9 amino acid residues of NeoE_4 was substituted by the 19 other natural amino acids was used (Fig. 6A). An HLA-A*02:01-positive lymphoblastic cell line was pulsed with these individual peptides and co-cultured with reporter cells (5KC NFAT ZsGreen) expressing TCR^127E10^. The level of activation of these reporter cells was assessed by cytometric detection of the fluorescence (Fig. 6B). The pattern of reactivity to this mutant library revealed the crucial and less permissive positions for activation (Fig 6C). The 42 single mutation peptides able to activate TCR^127E10^ do not occur in the annotated human proteome. We thus designed a combinatorial matrix that includes all amino acids positions enabling TCR^127E10^ activation. This peptide reactivity pattern (totaling 685440 possible peptides) was then searched in Uniprot which only yielded seven matches (Fig. 6D). These seven peptides were tested for activation of the reporter T cells. Only one peptide led to a detectable activation, but only with concentrations roughly 1000-fold higher than NeoE_4 (Fig. 6E). Finally, to investigate potential allogeneic activation, the reporter T cells were co-cultured with a panel of 21 lymphoblastic cell lines with diverse HLA alleles (Supplementary table S6). Notably, no allogeneic activation of TCR^127E10^ was observed in the presence of any of these cell lines unless loaded with exogenous NeoE_4 peptide (Fig. 6F). Altogether, these data indicate that neither off target nor allogeneic activation can be anticipated with TCR^127E10^-T cells thus highlighting the outstanding clinical potential of this TCR-T.

**Fig. 6:**
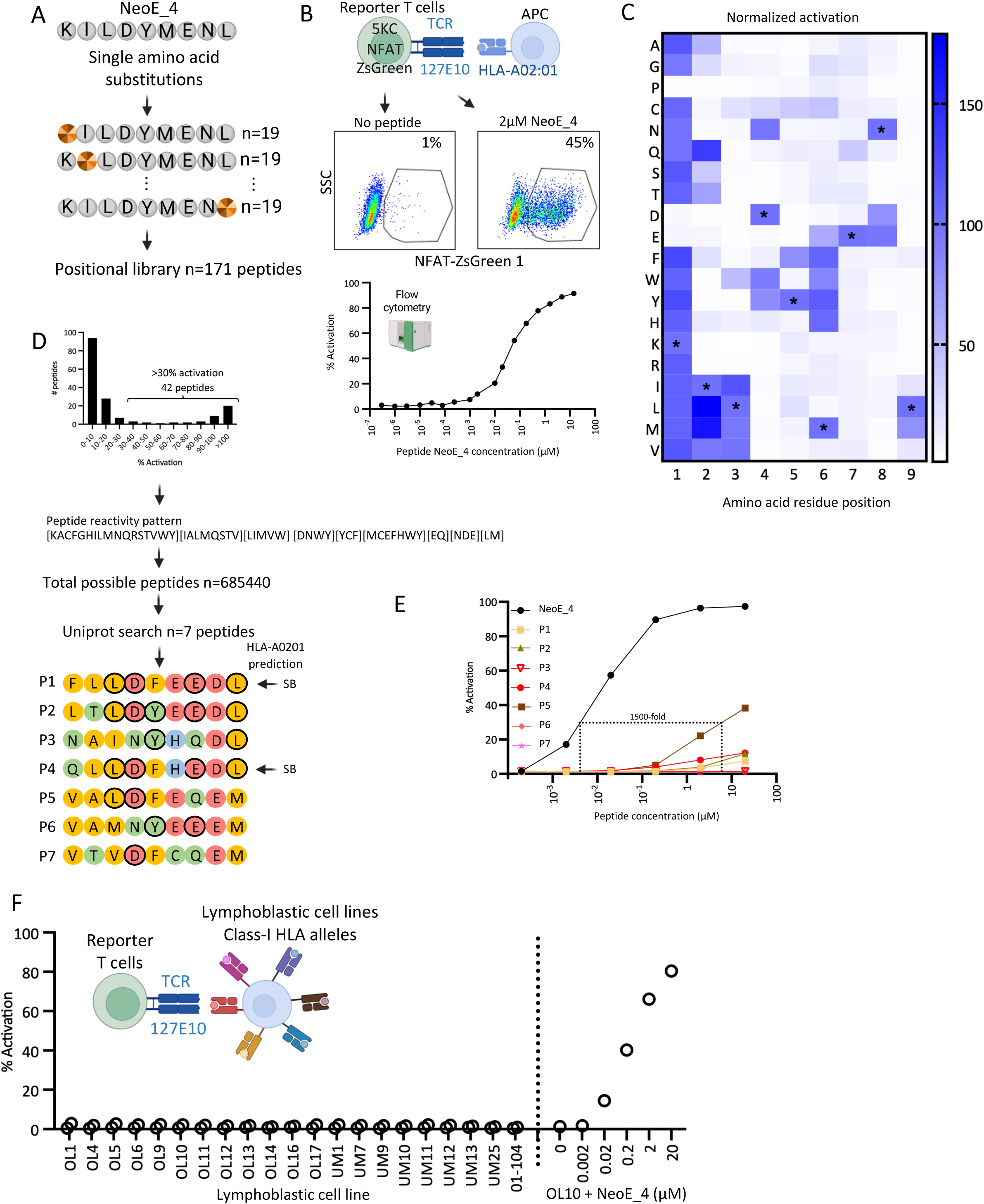
Safety of TCR127E10. (A). Peptide fingerprinting library of 171 peptides, each containing a single amino acid substitution of the target peptide NeoE_4. (B). Development of a standardized TCR activation assay using TCR127E10 transduced reporter T cells and flow cytometry. (C). Positional scan matrix for TCR127E10. Heatmap colors indicate the percentage of T cell activation in response to each variant normalized to NeoE_4 peptide. (D). From top to bottom, distribution of normalized activation induced by the 171 peptides, peptide reactivity pattern obtained with the 30% activation threshold, seven peptides which matches in the Uniprot database. Amino acid residue color code: yellow (hydrophobic), red (negative charge), blue (positive charge), green (hydrophilic uncharged). Black circles: residues shared with NeoE_4. (E), TCR activation assay with the seven peptides potentially cross-reactive and the NeoE-4. (F) Evaluation of TCR127E10 alloreactivity using reporter cells and a panel of 21 lymphoblastic cell lines. Cartoons created using Biorender.

## Discussion

Recent evidence from ribosome profiling and proteomic experiments suggests that peptides encoded by non-canonical ORFs (ORFs that are not reported in protein-coding sequence database), may encode microproteins with important biological functions in both disease and normal physiology (15–18). These non-canonical ORFs may be derived from alternative reading frames of protein-coding genes but also from long non-coding RNA (lncRNA) that may therefore not be as non-coding as their name suggests and encode microproteins with important biological functions. The estimated abundance of such noncanonical ORFs in the genome is a matter of intense debate (15).

The role of such microproteins in cancer is emerging. A growing number of reports indicate that HLA-I-bound peptides, derived from such non-canonical ORFs, constitute a significant fraction of the immunopeptidome raising a strong interest for immunotherapy approaches given the tumor-specific expression of some of these ORFs (19–22). However, to the best of our knowledge a clear demonstration that the unconventional immunopeptidome may constitute a target for immunotherapy has not been provided.

We recently discovered a novel class of lncRNAs that are uniquely regulated by oncogenic transcription factors in different sarcomas (9). In Ewing sarcoma, these lncRNA, also termed EwS*-*specific neogenes (*Ew_NG*s) are regulated by direct binding of EWSR1::FLI1, or related EwS-ETS fusion oncogenes, to microsatellite sequences leading to subsequent activation of transcription of intergenic regions. Their exclusive regulation by the disease-specific chimeric transcription factor gives them exquisite tumor specificity, as none of the normal tissues express *EWSR1::FLI1.* Thi*s* makes them, if translated and presented by HLA-I complexes, extremely attractive candidates for immunotherapy.

Here, we further investigated Ewing sarcoma cell lines and PDXs by riboseq and immunopeptidomics. In this study, we employed multiple layers of validation. Riboseq proved to be a valuable tool for prioritizing candidate neogenes, with *Ew_NG8* being the most robustly detected by this approach. Immunopeptidomics further corroborated *Ew_NG8* as a strong candidate through the detection of eight different HLA-I-bound peptides encoded by this neogene. T cells isolated from HLA-A*02:01-positive healthy donor, specific for a peptide derived from *Ew_NG8,* demonstrated, through both cell activation and functional assays, the specific detection of the peptide on the surface of EwS cell with the same HLA-I restriction. Strikingly, this T cell clone could kill all HLA-A*02:01-positive EwS cell lines tested, even those where the corresponding peptide was presented at low levels as it was undetected by immunopeptidomics. This indicates that T cell recognition is more sensitive in peptide detection than immunopeptidomics.

In addition to the HLA-I restriction, the killing was strictly dependent upon the expression of *EWSR1::FLI1* and of the source *Ew_NG8*. In addition, this cytotoxic property was fully reconstituted by transduction of the TCR into recipient T cells. These results demonstrate the ability of these *Ew_NG*-encoded peptides to be naturally processed in tumor cells as well as to be presented to T cells by HLA-I on Ewing cells and their potential to functionally activate T cells.

The presence of exquisitely Ewing specific neo-peptides presented by HLA molecules at the surface of EwS cells opens multiple therapeutic possibilities. Importantly, these peptides represent shared epitopes across EwS patients with a given HLA-I restriction. Therapeutic vaccination targeting different ORFs encoded by the *Ew_NGs,* following similar schemes developed for mutation-associated neopeptides would be an interesting option (23) facilitated by the shared nature of the antigens across patients. Bi-specific T cell engagers targeting EwS specific HLA-I peptide complexes also represent a potential strategy. Such a bi-specific reagent (tebentafusp) was recently successfully developed for the treatment of uveal melanoma (24). Finally, adoptive TCR-T cell therapy with autologous T cells genetically engineered to express a TCR able to recognize *Ew_NG-*derived antigens is an obvious very promising option, especially in the context of the recent FDA approval of the first TCR-T cell therapy, which targets a MAGE-A4 peptide presented by HLA-A*02:01 in aggressive synovial sarcoma (24). TCR-T^127E10^ identified here is a strong candidate for such a cell therapy as it is shown to have a strong activity *in vivo*. Moreover, our in depth analysis using a peptide scan approach and a search for allogenic activation demonstrated a favorable toxicity profilei. We are presently designing a first-in-class, first-in-child early phase clinical trial with this TCR-T.

Finally, our analysis of diverse tumors with oncogenic chimeric transcription factors (OCTF) suggests that neomorphic properties of these OCTF may also lead to transcription and potential translation of tumor-specific neogenes in other tumor types as well (9). If confirmed that these tumor present NG-encoded antigens, the immunotherapies that are highlighted above for Ewing sarcoma may be adapted to these other malignancies as well.

## Materials and methods

### Cell lines and patient-derived xenografts (PDXs)

A673 and A673/TR/shEF1 clone (also called ASP14) cell lines were cultured at 37°C, in 5% CO₂ with Dulbecco’s Modified Eagle Medium (DMEM) with High Glucose, 4 mM of L-Glutamine (Gibco), 4,500 mg/L Glucose and sodium 5-pyruvate (HyClone) supplemented with 10% FBS (Eurobio) and 1% antibiotics (v/v) (penicillin and streptomycin (Gibco). Blasticidin (10 μg/mL; Merk, Darmstadt, Germany) and Zeocin (100 μg/mL; Invitrogen, Waltham, MA, USA) were added to A673/TR/EF1 at 37 °C with 5% CO2. TC71, EW7, EW16, STA-ET-1, POE, MHH-ES1, Mel202, MP41 and MON cell lines were cultured in Roswell Park Memorial Institute (RPMI) 1640 medium (Sigma), 10% FBS (Eurobio), 1% antibiotics (v/v) (penicillin and streptomycin (Gibco). Culture cells were tested monthly for mycoplasma contamination.

EW7 cells were transduced with a lentiviral vector encoding a firefly luciferase–GFP (fluc-GFP) fusion reporter (Ew7-Luc). GFP-positive cells were isolated by fluorescence-activated cell sorting (FACS), clonally expanded, and subsequently assessed for stable GFP expression by flow cytometry several days after sorting. EwS patient-derived xenograft (PDX) models (IC-pPDX-5, IC-pPDX-8, IC-pPDX-141, IC-pPDX-164 and IC-pPDX-179) were generated at Institut Curie from patients under an Institutional-Review-Board-approved protocol (OBS170323CPP ref3272; dossier No. 2015-A00464-45). All PDX tumors exhibited a fusion transcript of *EWSR1::FLI1* except for IC-pPDX-164, showing *EWSR1::FEV* fusion (Supplementary Table S2).

### *EWSR1::FLI1* downregulation and CRISPR interference (CRISPRi)

Inhibition of *EWSR1::FLI1* in the A673/TR/shEF1 clone (ASP14) was induced by the expression of *EWSR1::FLI1* specific shRNA by adding 1 μg/mL of doxycycline in the medium ex-tempo. Specific *Ew_NG8* inhibition was obtained by CRISPRi method targeting the regulatory *EWSR1::FLI1*-bound microsatellite as previously described (9). Cells were transfected using lipofectamine RNAiMAX transfection Reagent (ThermoFisher) an equimolar mix of *Ew_NG8* or control guide RNAs and trackRNA (IDT) at a final concentration of 1 µM, diluted in OptiMeM medium (Gibco), for four days. Efficacy of CRISPRi was assessed by RT-qPCR.

### RT-qPCR

RNA was extracted with the RNeasy Plus Mini Kit (Qiagen) and reverse-transcribed using the High-Capacity cDNA Reverse Transcription Kit (Applied Biosystems). qRT-PCRs were performed using PowerSYBR green Mastermix (Applied Biosystems). Oligonucleotides were purchased from Eurofins Genomics. Reactions were run on CFX384 Touch Real-Time PCR instrument (Bio-Rad) and analyzed using the CFX Manager Software (Bio-Rad). Relative expression level was assessed with the ΔΔCt method using *RPLP0* as a housekeeping gene. Primer sequences are reported in Supplementary Table S4.

### Ribosome profiling

Ribosome profiling on 3 EwS cell lines (EW16, STA-ET-1 and TC71) and 5 EwS PDXs (IC-pPDX-5, IC-pPDX-8, IC-pPDX-141, IC-pPDX-164 and IC-pPDX-179) was performed by EIRNA Bio (https://eirnabio.com/). Briefly, samples were flash frozen and transferred into ice-cold polysome isolation buffer supplemented with cycloheximide (Millipore Sigma, #C1988). The library was sequenced using NovaSeq 6000 (Illumina). The per base sequencing quality of each three independent replicas passed the quality threshold. Ribo-seq data analysis was conducted using RiboTricer, including filter fastq files by quality, followed by removal of the adaptor sequence, alignment with GENCODE v19 reference transcripts and annotation of the *Ew_NGs*, using a minimal ORF length of 30 nucleotides. Translating ORF predictions were then performed using default parameters on Ribotricer QC data.

### Immunopeptidomics

Five Ewing sarcoma cell lines (A673, EW7, EW16, STA-ET-1, TC71) and five PDX (IC-pPDX-5,-8,-179,-164,-141) were used to search peptides associated to HLA-ABC. Cell suspensions were lysed in 50 mM Tris-HCl pH 8, 150 mM NaCl, 1X complete protease inhibitor (Roche), 5 mM EDTA, and 1% n-dodecyl-β-D-maltoside (Thermo Fisher Scientific # 89902), sonicated twice and incubated for 30 minutes at 4 °C with rotation. Samples were centrifuged at 20000 g for 1 hour at 4°C to remove insoluble fraction. Supernatants containing stabilized MHC-I-peptide complexes were quantified by BCA and incubated for 18h with CNBr activated Sepharose beads (GE Healthcare Life Sciences) coupled to anti–HLA-ABC W6/32 antibody (Bio X Cell). After immunoprecipitation samples were washed (i) three times with 50 mM Tris-HCl (pH 8.0), 150 mM NaCl, and 0.5% n-dodecyl-β-D-maltoside, (ii) three times with 50 mM Tris-HCl pH 8.0, 150 mM NaCl, (iii) one time with high salt concentration (50 mM Tris-HCl pH 8.0 and 0.5 M NaCl, (iv) three times with 50 mM Tris-HCl pH 8.0, 150 mM NaCl, (v) one time with 20 mM Tris-HCl pH 8.0. Then MHC-I-peptide complexes were eluted with 0.25% trifluoroacetic acid (TFA).

Samples were evaporated and resuspended in 30% acetonitrile (ACN), 0.1% formic acid (FA) and injected into an HPLC system (Agilent HP1100) for fractionation by strong cation exchange using a PolyLC sulfoethyl A column. Fractions were evaporated to dryness and reconstituted in 5% methanol / 0.5% TFA for mass spectrometry (MS) analysis. Samples were analyzed by liquid chromatography coupled to mass spectrometry (LC-MS/MS) using the high-resolution mass spectrometer 480 Exploris (Thermo Scientific) coupled to a Thermo Scientific Dionex Ultimate 3,000 chromatographic system. The separation was done at 300 nL/min, gradient 0-30% ACN (0.1% FA) in 60 min. The spectrometric analysis was performed in a data dependent mode and HCD fragmentation. The range acquired was 350-900 m/z.

LC-MS/MS spectra were searched using ProteomeDiscoverer 2.5 (ThermoFisher) and MSFragger (25) against predictive EwS neoprotein database (4,938 *Ew_NG* ORFs) (9), EwS fusion protein (6 fusions proteins) and Swissprot Human Reference Proteome. Peptides matching with annotated proteins were discarded. FDR was set to 1. HLA compliance between class I HLA alleles of the sample and amino acid motifs of the peptides was predicted using NetMHCpan 4.0.

### Class I HLA-peptide binding assays and tetramers

Biotinylated recombinant class I HLA molecules were purchased from immunAware (Copenhagen, Denmark) as easYmers (catalog #1002-1). Peptides were synthetized at >95% purity (GeneCust). Monomers were loaded with peptides by incubation at 18 °C for 48 hours (easYmers). Binding of peptide to class I HLA was measured by flow cytometry following the manufacturer’s instructions. HLA/peptide monomers were incubated with streptavidin-coated beads, stained with anti-human b2-microglobulin antibody coupled to phycoerythrin and analyzed by flow cytometry. Peptides used as positive controls (100%) of HLA binding were provided by the manufacturer: HLA-A*01:01 (YFV NS5 286-295, KSEYMTSWFY), HLA-A*02:01 (CMVpp65 495-503, NLVPMVATV), HLA-A*03:01 (CMV IE1 99-107, RIKEHMLKK), HLA-A*68:01 (CMV IE1 33-41, TTFLQTMLR), HLA-B*07:02 (CMV pp65 417-426,

TPRVTGGGAM), HLA-B*51:01 (Cryptosporidium parvum, VPFVSVNPI), HLA-B*57:01 (YFV NSI 116-124, KTWGKNLVF). For each tetramer, HLA/peptide complex (100 μM) was combined for 1 hour at room temperature with fluorescent streptavidin (BioLegend). Tetramers were stored at 4 °C for a maximum of 3 months.

### Tetramer & antibody staining

Peripheral blood mononuclear cells (PBMCs) were isolated using standard Ficoll-gradient procedures and either studied fresh for phenotyping or frozen. PBMC were thawed in RPMI medium (GIBCO) containing 10% FBS. Cell suspensions were incubated for 30 minutes in culture medium containing 50 nmol/L dasatinib (Lissina et al., 2009) to improve tetramer staining. To isolate and clone CD8^+^ T cells specific for a given HLA-peptide, healthy donors bearing the corresponding HLA allele were selected. 2×10^8^ PBMC were first stained with live/dead aqua (Invitrogen) followed by the peptide-loaded tetramers associated to PE and APC for 20 minutes at room temperature. Then cells were washed and incubated with a mix of anti-PE and anti-APC microbeads (Miltenyi) for 20 minutes at 4°C. After washing in PBS+ (PBS 1% human male AB serum (Biowest), 2 mM EDTA (Invitrogen)) tetramer positive cells were enriched in a MS column (Miltenyi). Enriched cells were stained with anti-CD45RA FITC (BD 566349), CCR7 BV421 (BioLegend 353208), CD8 BUV395 (BD 563795) and CD3 BUV737 (BD 612750) for 20 minutes at 4°C. Cells were then washed and analyzed in an ARIA Fusion cell sorter (BD). Data were analyzed using FlowJo V10.10 software (BD).

### T cell clone generation

Double tetramer positive CD8^+^ single cells were sorted into 96 well plates containing 1:1 AIM-V/RPMI medium supplemented with 5% human serum, 100 U/mL penicillin, 100 μg/mL streptomycin in the presence of 2×10^5^ irradiated (25 Gy) feeder cells. Cells were stimulated with 300UI/ml human IL-2 (Proleukin, Novartis) and 30 ng/ml anti-CD3 (OKT3, eBioscience). Growing clones were cultured for 3 weeks in the same medium without addition of anti-CD3 and tested for tetramer staining. Tetramer positive clones were stimulated every 3 weeks using the same media containing IL-2, anti-CD3 and irradiated feeders. After each cycle of amplification, the clones were tested for tetramer binding by cytometry. cDNA from each clone was amplified by PCR using primers for TRAV, TRBV, and constant regions (26), the PCR products sequenced and the resulting sequences analyzed using IMGT/V-QUEST (27).

### T cell clone activation

The avidity of the clones for HLA-A2 restricted peptides was assessed after coculture of T cell clones with K562-HLA-A*02:01 cell line loaded with peptide. Briefly, the cells were washed in RPMI without serum and incubated for 2 hours at 37°C at increasing doses of peptide (0.3 pM to 14 µM), then washed and co-cultured with clone cells at 1:1 ratio for 18 hours. The supernatants were collected and the secreted IFN-γ, TNF-α, Granzyme B and Perforin quantified using LEGENDplex (BioLegend) and a Cytoflex cytometer (Beckman).

### TCR-T cell generation

To generate TCR-T cells bearing the 127E10 TCR specific for neoE_4 peptide, the TCR alpha and beta chains (Supplementary Table S5) were cloned into a lentiviral vector bearing murinized TRA and TRB constant regions (13). The same was performed for a CMV pp65 (NLVPMVATV) specific TCR (Supplementary Table S5). Lentivirus were prepared by VectorBuilder. Human CD8^+^ T cells from HLA-A*02:01 negative healthy donors were isolated from PBMC using human CD8^+^ T cell enrichment kit (StemCell #19053) and frozen or used fresh. CD8^+^ T cells were activated for 2 days in Xvivo 15 (Lonza) media supplemented with 5% human AB serum, 50 µM b-mercaptoethanol, 300 UI/ml IL-2, 5 ng/ml IL-7, 5 ng/ml IL-15 and Dynabeads human T-activator CD3/CD28. The cells were then transfected with the TCR encoding lentivirus at MOI 5 in the presence of polybrene 5 ng/ml. Lentivirus were spinoculated for 45 minutes at 1000 g and 32°C. At day 6 the Dynabeads were removed and the exogenous TCR expression tested by flow cytometry using anti-mouse TCRβ BV711 (BioLegend 109243), anti-human CD8 BUV395 (BD 563795) and CD3 BUV737 (BD 612751). When indicated, IFN-γ and TNF-α release was detected using the kit ELISA MAX Deluxe (BioLegend), following the manufacturer’s instructions.

### *In vitro* tumor cell killing

The capacity of the cloned T cells and engineered TCR-T cells to kill Ewing sarcoma cell lines was assessed using a killing assay in an IncuCyte system. Tumors cells were plated in 96-well plates (50,000 cells per well) and cultured overnight to allow cell adhesion. RPMI media was complemented with 1 mM CaCl2. Then, Annexin V Red dye (Sartorious) was added following manufacturer’s instructions. As a control, in some wells the interaction between TCR and peptide MHC was blocked by the addition of anti-HLA (W6/32) antibody (10 µM). Effector T cells (clone or TCR-T) were added at different effector:target (E/T) ratios. As a positive control the cognate peptides (1 µM) were added. Cell killing was monitored and analyzed by an IncuCyte ZOOM live cell analysis system (Sartorius) for 24 hours. Four images per well at 40x zoom were collected at each time point. Tumor growth was quantified by measuring total integrated Annexin V Red intensity per well every 4h. Annexin V Red signal was normalized to the time 0 signal. Supernatants were collected at 24 hours, and TNF-α and IFN-γ were determined by ELISA as above. Duplicates were plated for each condition, and t-test analysis was applied.

### Luciferase-based cytotoxic assay

5 × 10^4^ EW7-Luc cells (target cells) were cocultured with CD8 T cells from day 0 (effector cells) in triplicates at different effector cells to target cells (E:T) ratios (5:1, 1:1, and 1:5) in 100 μL/well of complete Xvivo using a U bottom 96-well plate. Target cells were plated at the same cell density alone to determine the maximal luciferase expression (relative light units: RLU). Effector cells were plated alone to measure the background. Eighteen hours after culturing the cells at 37°C, 50 μL of luciferase substrate at 0.3 mg/mL in PBS was added to each well, and luminescence was detected in a SpectraMax ID3 plate reader (Molecular Devices). % Specific lysis was determined as [1 − (RLUsample − RLUbackground)/(RLUmax − RLUbackground)] × 100. Untransduced T cells cells were used as effector negative control.

### xCELLigence cytotoxicity assay

A real-time, impedance-based cytotoxicity assay was performed using the xCELLigence Real-Time Cell Analyzer (RTCA) eSIGHT system (Agilent) to assess the killing of HLA-A02:01 EW7–Luciferase target cells. 2.5×10^3^/well HLA-A*02:01 EW7–Luciferase cells were seeded in collagen-precoated 96-well E-plates. After 24 h, CD8⁺ T cells expressing TCR127E10 were added at effector-to-target (E:T) ratios of 5:1, 1:1, and 1:5. Cell impedance (reported as Cell Index) was monitored every 15 min for 48 h. Background impedance (blank) was measured in wells containing complete medium alone. Data were normalized to the Cell Index value at the time of effector cell addition. Untransduced CD8⁺ T cells and tumor cells alone were included as effector-negative and target-negative controls, respectively.

### Flow Cytometry

TCR-T cells activation by tumor cells was assessed using CD137 expression measured by flow cytometry. Tumors cells were plated in 96-well plates (density of 50,000 cells per well in 100ul of media) and cultured overnight to allow cell adhesion. Effector T cells (TCR-T or unstransduced) were added at an effector:target (E/T) ratio of 1:1 and cultured for 24h. As control, cultured TCR-T cells without tumor cells were used. Then TCR-T cells were stained with various antibodies using anti-CD8 BUV395 (BD563795), anti-CD3 BUV 737 (BD 612751), anti-mTCR BV711 (Biolegend 109243) and anti-4-1BB BV650 (Biolegend 309828) for 20 minutes at 4°C. All antibodies were diluted using the BD Horizon Brilliant Stain Buffer (BD Bioscience). Samples acquisition was performed with the BD LSRFortessa™ and FACSDiva software. Results were generated using FlowJo software (BD) and Graphpad Prism 10.

### Safety assessment of TCR127E10 by peptide fingerprinting

A positional scanning library (14) of 172 9-mer peptides encompassing all possible single amino acid residue substitutions in NeoE_4 peptide was generated. EBV-OL-10, an HLA-A*02:01 lymphoblastic cell line was pulsed with individual peptides and co-cultured with 5KC cells bearing the NFAT promoter-ZsGreen-1 reporter (28) and transduced with TCR127E10. Activation was assessed by flow cytometry. To calibrate the assay, we used antigen presenting cells pulsed with serial dilutions of the target peptide and quantified reporter cell activation. For the cross-reactivity tests, peptides were incubated at 0.2 and 2 µM. Results were normalized to activation with the NeoE_4 peptide. The activation threshold was determined as 30%. Substitutions above this threshold were retained to construct the positional matrix. The 685440 possible peptides from the matrix were searched in Uniprot. The seven matches (FLLDFEEDL, LTLDYEEDL, NAINYHQDL, QLLDFHEDL, VALDFEQEM, VAMNYEEEM, VTVDFCQEM) were synthetized and subjected to the activation test. Allogeneic activation was tested using a panel of 21 lymphoblastic cell lines encompassing 13 HLA-A, 16 HLA-B and 16 HLA-C alleles (Supplementary Table S6).

### In vivo tumor model and treatment

To evaluate the in vivo efficacy of CD8⁺ T cells expressing TCR^127E10^, 1×10⁶ HLA-A*02:01 EW7-Luciferase cells were injected intraperitoneally (i.p.) into NSG mice. Seven days after tumor engraftment, mice received an intraperitoneal injection of either 1×10⁷ human CD8⁺ TCR^127E10^-positive T cells or untransduced CD8⁺ T cells, depending on the experimental group. Recombinant human IL-2 (2,500 IU) was administered daily by intraperitoneal injection from the day of T-cell transfer until day 25. Tumor burden was monitored twice weekly by bioluminescence imaging using the Xenogen IVIS Imaging System (PerkinElmer). Bioluminescence data were analyzed with Living Image software (PerkinElmer) and are presented as average radiance (photons/s/cm²/sr). Humane endpoints were defined as an average radiance exceeding 1×10⁷ photons/s/cm²/sr, the presence of combined clinical signs of distress, or a maximum body weight loss of 20%. The experiment was conducted with 12-week-old NOD/SCID/IL-2Rγ-null (NSG) female mice purchased from Charles River Laboratories (France), in an accredited animal facility by the French Veterinarian Department following ethical guidelines, approved by the relevant ethical committee (APAFIS##47660-2024020113137204 v4, DAP 2017-023).

## List of Supplementary Materials

Supplementary Fig. S1. Immunopeptidomic data

Supplementary Fig. S2. Riboseq data

Supplementary Figure S3. HLA-A*02:01 expression and *EWSR1::FLI1* or *Ew_NG8* invalidation

Supplementary Figure S4. TCR-T cells activation by EwS and non EwS cell lines

Supplementary Figure S5. Characterization of TCR^127E10^ for *in vivo* experiments

Supplementary Table S1. Summary of identified peptides and proteins per sample

Supplementary Table S2. Genetic and HLA characterization of EwS samples

Supplementary Table S3. Ew neogenes and *EwS-NG* ORFs encoding for EwS neopeptides

Supplementary Table S4. Primer sequences used for qPCR

Supplementary Table S5: TCR sequences.

Supplementary Table S6: HLA restriction of lymphoblastic cell lines used to test TCR127E10 alloreactivity

## Acknowledgments

We thank all members of the Diversity and Plasticity of Pediatric Tumors, the Innate-like and CD4 T cells in cancer and Integrative Functional Genomics of Cancer laboratories for helpful discussions. We also thank the Cytometry and Imaging platforms of institute Curie for outstanding assistance.

We are grateful to Francesca Lucibello, Abdoulaye Soumare, Christina Ekwegbara, Stéphanie Fitte-Duval, Giacomo Oliveira, Pierre Fumeron, Olivier Ganier, Céline Chauvin, Pierre-Grégoire Coulon, Joanna Cyrta for advice, discussion and assistance.

This work was supported by Institut Curie, Institut national de la santé et de la recherche médicale (INSERM 23CP082-00), Ligue Nationale Contre le Cancer (AAPEAC2020.LCC/OD), Institut National du Cancer (PLBIO19-192, PLBIO19-140, high risk high gain 2023-22), Agence Nationale de la Recherche (ANR-10-EQPX-03, ANR-20-CE19-0012-03), Institut Curie Génomique d’Excellence (ICGex), Société Française de lutte Contre les leucémies et cancers de l’Enfant et de l’adolescent, Janssen Horizon (CT9332), Fight Kids Cancer (21-FKC-EOI-038).

This project also received support from European funding as follows: ERA-NET TRANSCAN JTC-2011 (01KT1310), TRANSCAN JTC-2014 (TRAN201501238), TRANSCAN JTC-2017 (TRANS201801292), EEC (HEALTH-F2-2013-602856), H2020-lMI2-JTl-201 5-07 (116064—ITCC P4), and H2020-SC1-DTH-2018-1 (SEP-210506374—iPC).

We are indebted to the following associations for providing essential support: la Course de l’Espoir, M la vie avec Lisa, ADAM, Couleur Jade, Dans les pas du Géant, Courir pour Mathieu, Marabout de Ficelle, Les Bagouzamanon, Enfants et Santé, Les Amis de Claire, Un Elan pour Lucas, and Amarape. J.V. and CC were supported by a Ligue Nationale Contre le Cancer and Institut Curie PhD fellowship. O.S. was supported by a fellowship from the French Ministry of Higher Education and Research and Institut Curie.

## Authors’ Contributions

**A. Lalanne:** Investigation, performed experiments, visualization, methodology, prepared figures, writing–original draft. **C. Collin:** Investigation, designed and performed experiments. **F. Petit:** Provided critical help with various experiments, analyzed data, prepared figures, and drafted the manuscript. **M. Lacaud:** Provided help with TCR-T related experiments, data analysis, prepared figures, in vivo experiments and EW7_Luc generation. **Y. A. Arribas:** Provided critical help with immunopeptidomic experiments and data analysis. **A. Darbois Delahousse**: Performed in vivo experiments. **A. Leruste:** Contributed conceptual advice. **M. Koshkina**: Assisted with bioinformatics analysis. **K. Raymond**: provided help with TCR-T related experiments. **P. Klein:** Assisted with bioinformatics analysis. **J. Vibert:** Assisted with bioinformatics analysis and developed the EwS-specific neoprotein database. **S. Zaidi:** PDXs data curation and provided help with coculture experiments. **S. Grossetete:** Contributed help on computational biology analyses**. J. Pilet:** Help on data analyses of neogenes. **K. Laud-Duval:** Advice on experimental design. **S. Aflaki:** Contributed with T cell clone culture, characterization, TCR sequencing, TCR re-expression and safety assays. **C. Jamet:** contributed with cell culture, TCR sequencing and TCR re-expression. **M. Carrascal:** Contributed with critical advice on immunopeptidomics analysis and performed spectra validations. **J. Maggi:** Contributed with critical advice on immunopeptidomics analysis and performed spectra validations**. S. Menegatti**: Provided conceptual advice. **J. Fuentealba**: Provided conceptual advice. **M. Alcantara**: Provided conceptual advice. **J. Waterfall:** Conceptualized the project, supervised bioinformatic work, funding acquisition, validation. **O. Lantz:** Conceptualization, data curation, formal analysis, supervision, funding acquisition, validation, writing–original draft and editing. **O. Delattre:** Conceptualization, data curation, formal analysis, supervision, funding acquisition, validation, writing–original draft and editing.

---

**Supplementary Fig. S1.**
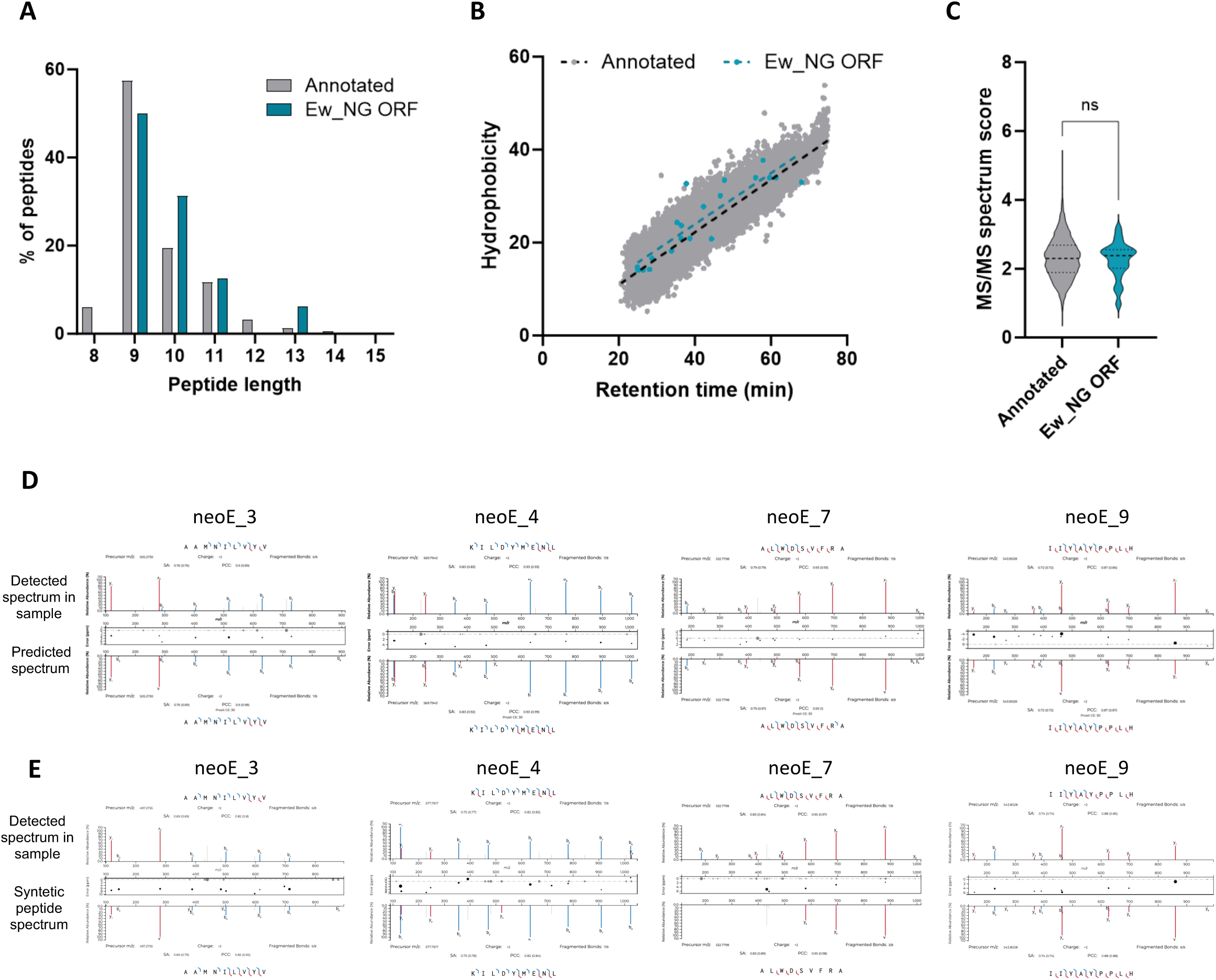
Immunopeptidomic data. **(A)** Peptide length distribution (in amino acids) from annotated and *Ew_NG*-derived peptidomes (n=16). **(B)** Scatter plots comparing retention time and hydrophobicity index from annotated and *Ew_NG*-derived peptides obtained with ProteomeDiscoverer. **(C)** Boxplot showing the MS/MS peptide-spectrum identification score (SEQUEST score) from annotated-canonical and *Ew_NG*-derived peptides. **(D)** Comparison of spectra from *EwS_NG*-derived peptides identified in EwS samples (top) with spectra predicted using Prosit (bottom). **(E)** Comparison of spectra from *EwS_NG*-derived peptides identified in EwS samples (top) with spectra of synthetic peptides (bottom). Spectrums were extracted from Proteome Discoverer and plotted using Universal Spectrum Explorer. A total of 16 EwS_NG-derived peptides have been validated after comparing the fragmentation patterns between endogenous and synthetic peptides.

**Supplementary Fig. S2.**
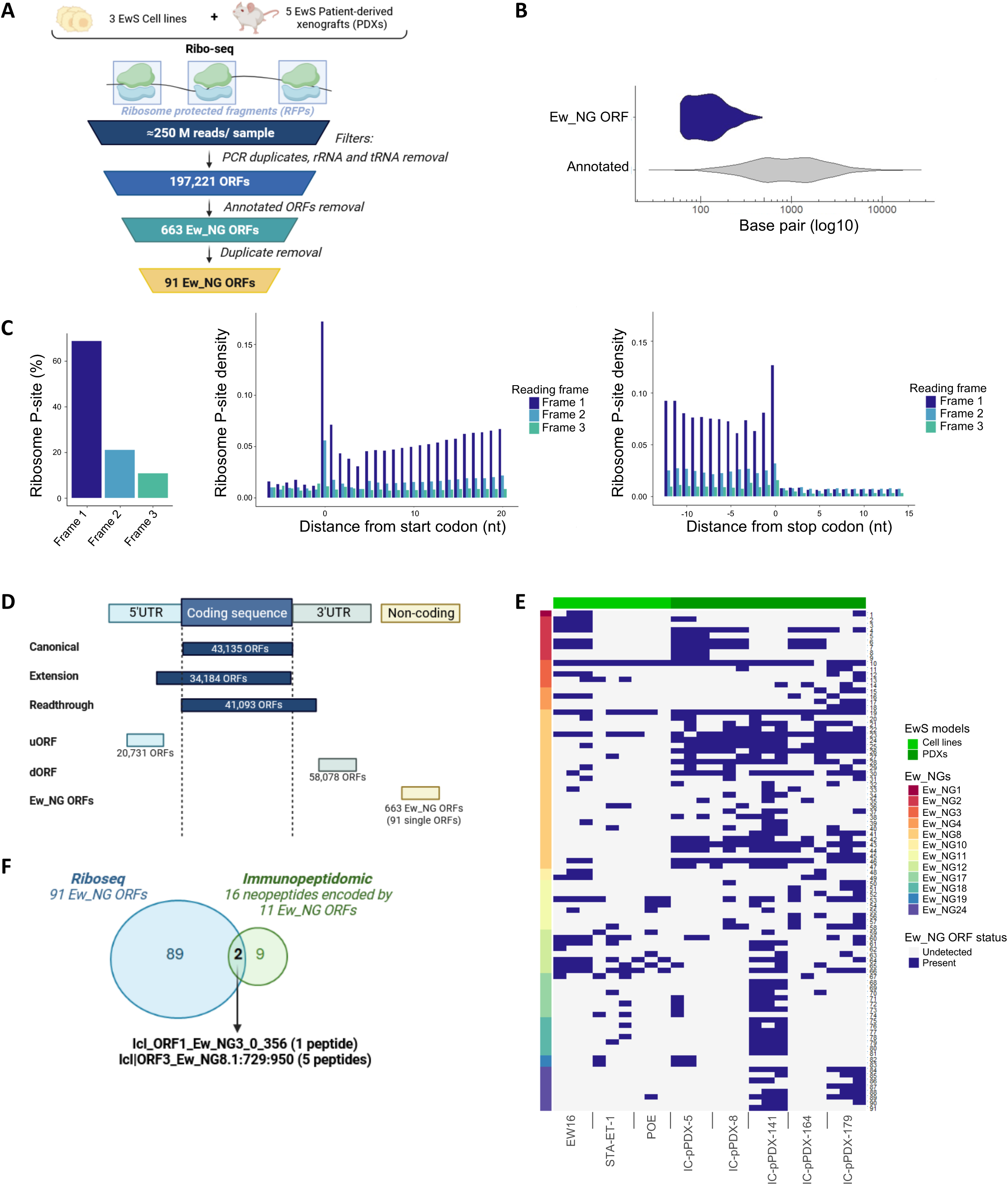
Riboseq data. **(A)** Overview of the experimental design for Riboseq analysis in EwS cell lines and PDXs (Created by Biorender). **(B)** Comparison of ORFs length between *Ew_NG*-related ORFs and canonical annotated ORFs. **(C)** Position of the Riboseq P-sites with respect of the three potential reading frames and bar plot of RPF density along the translation start and end sites in the EW16 EwS cell line **(D)** Different types of ORFs detected **(E)** Heatmap summarizing Ew_NG ORFs found in each Riboseq sample analyzed. **(F)** Overlap between *Ew_NGs*-derived RPFs and immunopeptides.

**Supplementary Figure S3.**
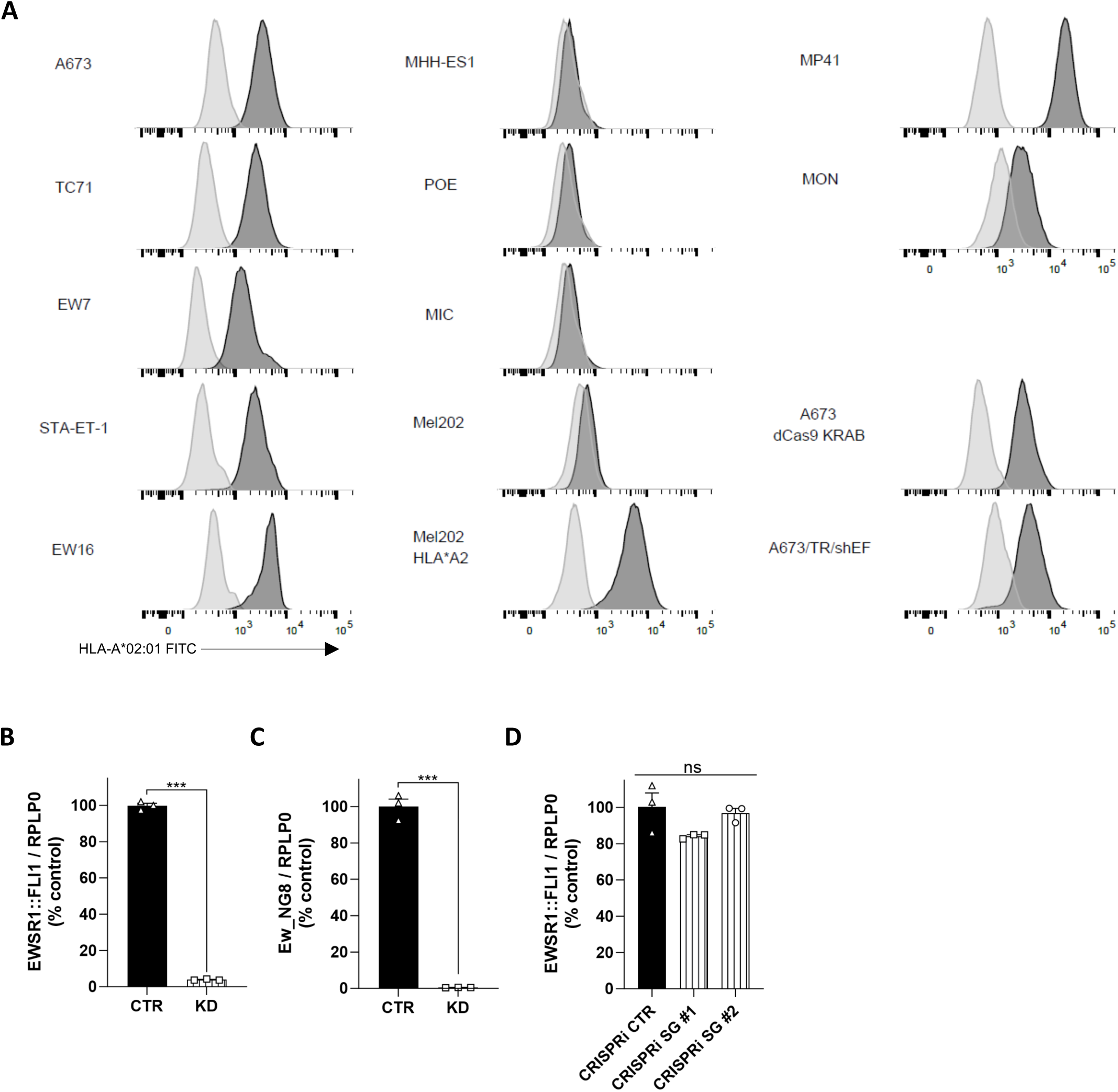
HLA-A02 expression and *EWSR1::FLI1* or *Ew_NG8 invalidation*. **(A)** Flow cytometry assessment of HLA-A02 expression in EwS or non EwS cells (black) compared to an isotype control (grey). **(B-C)** RT-qPCR of *EWSR1::FLI1* (B) or *Ew_NG8* expression (C) in the DOX-inducible A673/TR/shEF EwS cells. CTR, cells without DOX; KD, cells incubated with DOX. Results are represented as mean ± SEM, *n=3*. **(D)** RT-qPCR of *EWSR1::FLI1* in the A673 EwS cells after CRISPR interference of *Ew_NG8* with control (CTR) or with two different *Ew_NG8*-specific single guides (SG #1 and #2). Results are represented as mean ± SEM, *n=3*. *** *p* value < 0.001. ns, non-significant.

**Supplementary Figure S4.**
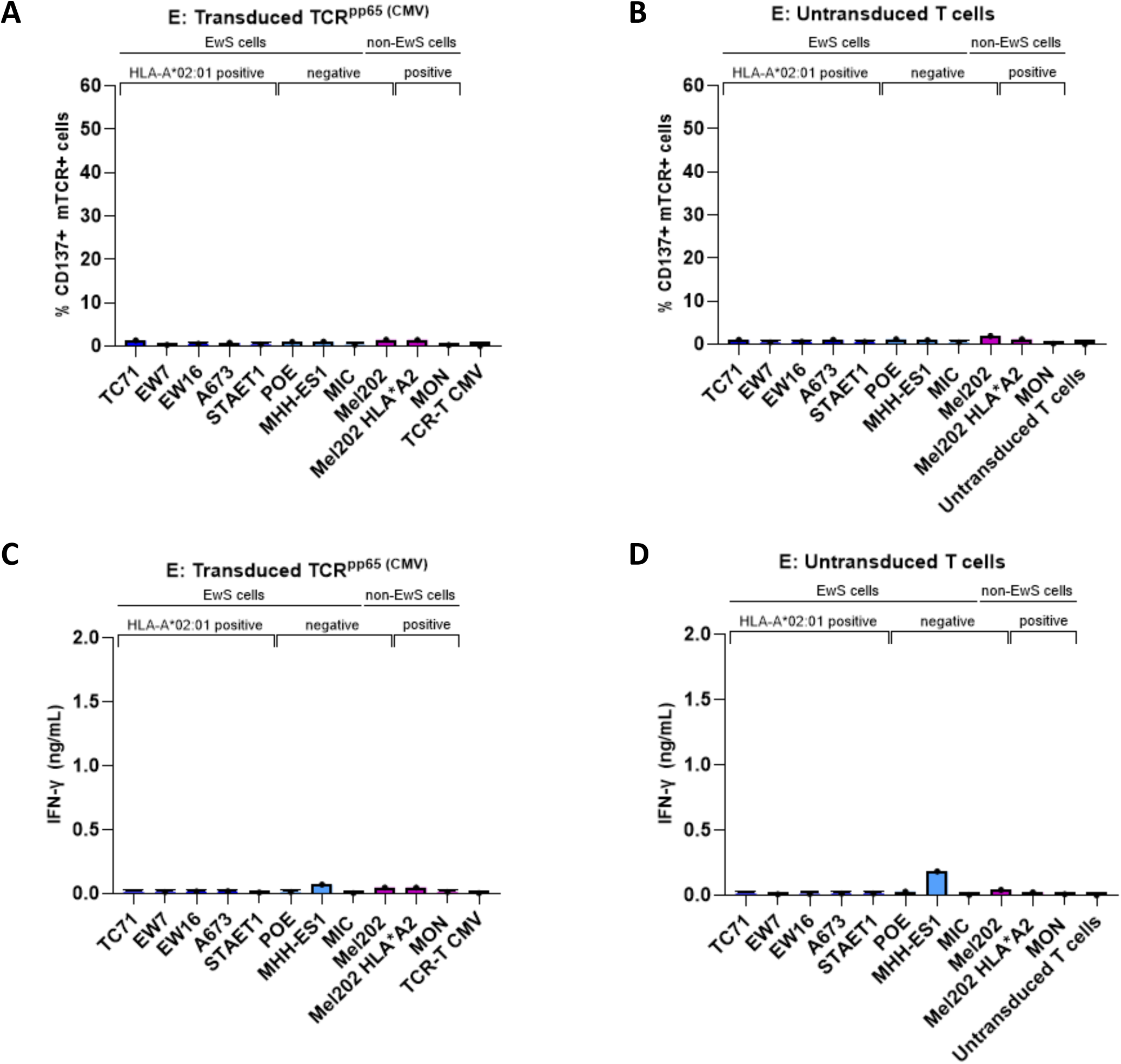
**TCR-T^pp65^ and untransduced T cells activation by EwS and non EwS cell lines and IFN-γ secretion (A-B**) %CD137^+^ mTCR^+^ of (A) transduced TCR^pp65^ T cells or (B) untransduced T cells after activation with EwS (blue) or non-EwS (purple) cell lines. **(C-D)** IFN-γ cytokine secretion by (C) transduced TCR^pp65^ T cells or (D) untransduced T cells after activation with EwS (blue) or non-EwS (purple) cell lines. Results are represented as mean ± SEM, *n=1*.

**Supplementary figure S5.**
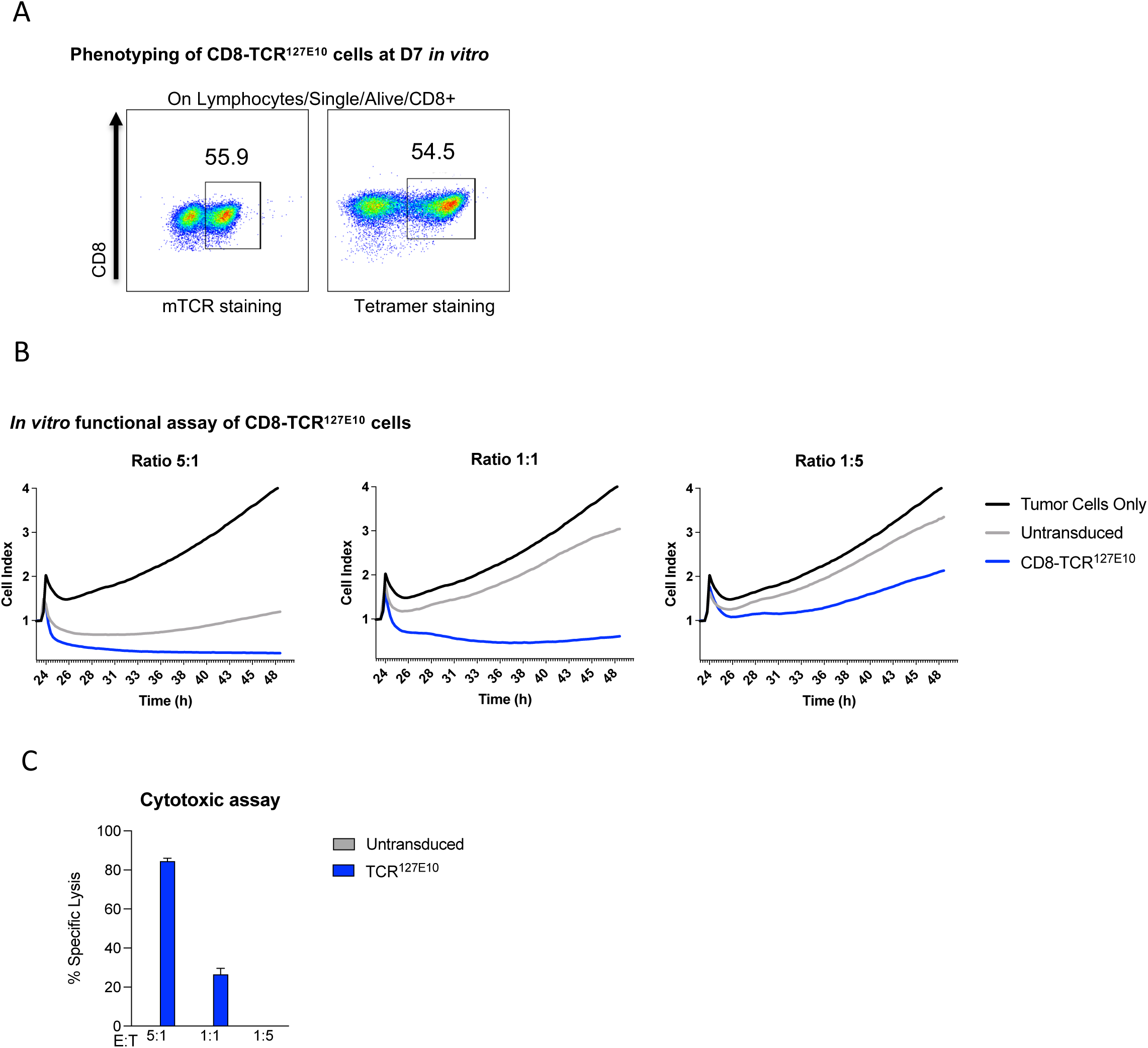
TCR-T cells used for in vivo assays. **(A)** Representative FACS plots and percentages of CD8-TCR^127E10^ measured 7 days post transduction with either an ⍺-murine TCR antibody or an HLA-A*02:01 peptide specific tetramer. **(B)** *In* vitro killing activity of CD8-TCR^127E10^ on HLA-A*02:01 Ew7-Luc cells by impedance mesurement. HLA-A*02:01 EW7-Luc cells were seeded (2.5 x 10^3^ per well in collagen-precoated 96-well E-plates, in triplicate). 24 h later, untransduced CD8 or CD8-TCR^127E10^ were added at effector-to-target ratios (E:T) of 5:1, 1:1, and 1:5. Cell impedance (reported as Cell Index) was monitored every 15 min for 48 h. Data were normalized to the Cell Index value at the time of effector cell addition. **(C)** Luciferase-based cytotoxicity assay showing CD8-TCR^127E10^ T cells or untransduced CD8 T cells co-cultured with EW7-Luc cells at different effector-to-target (E:T) ratios (left panel) for 24 h. Bars represent the mean of triplicate wells.

